# Mechanical Cues Regulate Estrogen and Progesterone-Induced Nascent ECM Deposition by Human Endometrial Stromal Cells

**DOI:** 10.1101/2025.10.14.682403

**Authors:** Grace K. Hinds, Arina Velieva, Yu-Chung Liu, Avinava Roy, Rima Chavali, Claudia Loebel

## Abstract

The endometrium, the mucosal lining of the uterus, is a highly regenerative tissue that undergoes cyclic remodeling guided by tightly regulated levels of estrogen and progesterone. Stromal cells are embedded within the connective tissue of the endometrium and contribute to the rapidly changing extracellular matrix (ECM). With hormone exposure, endometrial stromal cells undergo decidualization, which alters their morphology and protein secretion. While an increase in tissue modulus is associated with gynecological diseases, the relationship between mechanical properties, hormone exposure, and ECM deposition remains poorly understood. Here, we investigated how both stiffness and hormones regulate ECM deposition by human endometrial stromal cells during decidualization. Using metabolic labeling with sugar analogs and click chemistry, we measure newly secreted ECM proteins deposited by endometrial stromal cells during decidualization. Additionally, we study the nascent ECM in response to different mechanical properties using hyaluronic acid hydrogels. To increase throughput and reproducibility, we designed an automated ImageJ-based workflow for unbiased quantification of nascent ECM deposition. Our results demonstrate that hormones induce decidualization, characterized by F-actin stress fiber formation and prolactin secretion. In addition, we show that decidualization on hydrogels is characterized by an increase in nascent ECM deposition which depends on the initial hydrogel modulus. In contrast, endometrial stromal cells on glass show little change in nascent ECM deposition during hormone exposure. Collectively, these findings demonstrate that both mechanical and biochemical cues regulate ECM deposition during endometrial remodeling. These observations may provide new insights towards future studies addressing the mechanisms of ECM remodeling in gynecological diseases.

## 1. Introduction

The endometrium is a highly regenerative tissue that, together with the myometrium (muscle layer), the fallopian tubes, and ovaries, forms the female reproductive system (Figure 1a). The endometrium consists of approximately 80% stromal cells that are embedded in a laminin-, collagen-, and fibronectin-rich extracellular matrix (ECM)^[1]^. Each month, the ovaries release controlled amounts of hormones, including estrogen and progesterone. These hormone levels are cyclical, with estrogen peaking at days 10-12 (prior to ovulation) and progesterone at days 17-21(prior to menstruation, Figure 1b). Guided by these hormonal fluctuations, the endometrium undergoes a complex and tightly regulated process of growth (proliferative phase), maturation (remodeling phase) and finally degradation (menstruation, Figure 1c). During monthly regeneration, the endometrial tissue grows up to 9 mm before it is remodeled and degraded during menstruation ^[2]^. This process occurs in a scar-free manner and repeats up to 400 times during a reproductive lifespan ^[3]^. Thus, it requires precise control over cellular activity and interactions with the ECM. Studying the core mechanisms of the complex endometrial functions offers potential to further understand tissue regeneration and cell-matrix interactions. Endometrial stromal cells are central to this cell-ECM crosstalk ^[4]^ and have been shown to contribute to ECM deposition and remodeling ^[5],[6]^.

**Figure 1.**
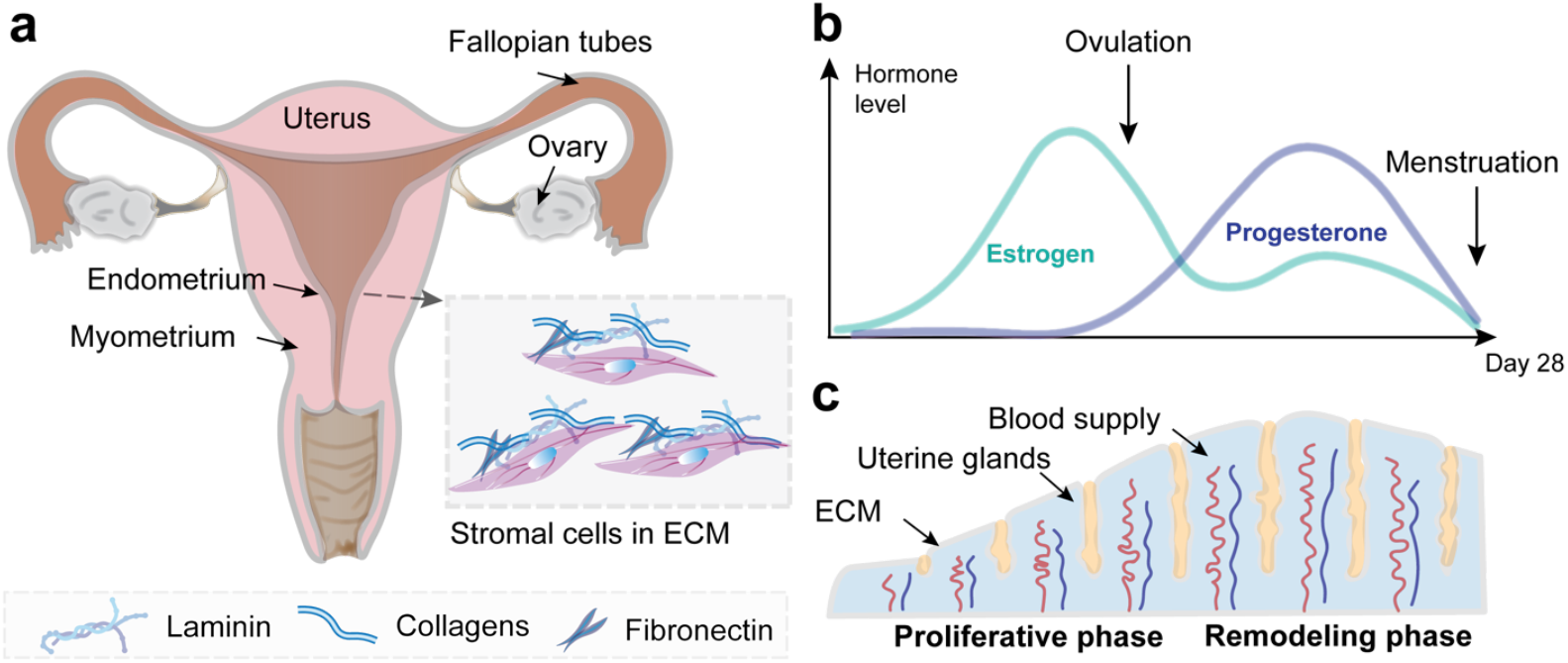
Endometrial tissue undergoes cyclic remodeling in response to estrogen and progesterone. **a** Schematic illustrating the female reproductive system including the uterus with the endometrium and myometrium, the ovary and fallopian tubes. Inset shows endometrial stromal cells embedded in extracellular matrix (ECM) containing a variety of proteins (e.g., laminin, collagens and fibronectin). **b** Schematic illustrating the relative levels of estrogen and progesterone during the menstrual cycle. **c** Schematic illustrating the endometrial tissue during the proliferative and remodeling phase of the menstrual cycle.

A critical part of endometrial remodeling is decidualization - during which endometrial stromal cells differentiate into decidual cells to prepare the endometrium for embryo reception. Successful decidualization has been associated with the controlled remodeling of ECM ^[7]^. In contrast, increased endometrial tissue modulus is characteristic of gynecological pathologies such as adenomyosis (growth of endometrial tissue into the myometrial layer) and endometrial cancer ^[8],[9]^. While previous work has qualitatively described changes to ECM density and arrangement throughout the menstrual cycle, how hormones and tissue stiffness direct ECM remodeling and deposition remains unknown ^[10]^.

To address this gap in knowledge, we employed our previously established metabolic labeling technique to measure newly secreted (nascent) ECM deposition by human endometrial stromal cells with and without hormones (estrogen and progesterone). In addition, we used hyaluronic acid hydrogels of varying elastic moduli to establish relationships between hormone levels, hydrogel modulus, and ECM deposition.

## 2. Experimental

### 2.1 Hydrogel synthesis and fabrication

#### Norbornene-modified hyaluronic acid (NorHA) synthesis

NorHA was synthesized as previously described ^[11]^. Briefly, sodium hyaluronate was dissolved in 2-(N-morpholino)ethanesulfonic acid (MES) buffer (pH 5.5) at 1% weight/volume (w/v). Next, 4-(4,6-Dimethoxy-1,3,5-triazin-2-yl)-4-methylmorpholinium chloride (DMTMM) and 5-norbornene-2-methylamine (Nor) were added sequentially at a 1:3:2 molar ratio (HA:DMTMM:Nor) and stirred for 24 hours at room temperature. This was followed by the addition of 8% (v/v) saturated sodium chloride and ethanol precipitation. The precipitate was collected by vacuum filtration, re-dissolved in deionized water, dialyzed against water for 72 hours (6 - 8 kDa molecular weight cut-off) and lyophilized. The degree of norbornene modification (21%) was confirmed by proton nuclear magnetic resonance (^1^H NMR).

#### Hydrogel fabrication

First, 12 mm circular cover slips were methacrylated through incubation in ethanol containing 0.3 v/v% glacial acetic and 0.5 v/v% 3-(Trimethoxysilyl)propyl methacrylate for 6 min at room temperature, followed by air-drying for 30 min. NorHA hydrogels were fabricated by mixing a 4 wt% pre-polymer solution with 0.05 wt% lithium phenyl-2,4,6-trimethylbenzoylphosphinate (Colorado Photopolymer Solutions), dithiothreitol (DTT), and 1 mM thiolated-RGD (Genscript). Pre-polymer solutions were pipetted as droplets onto Sigmacote-treated rectangular cover glasses and a methacrylated 12 mm circular cover glass was placed on top of the droplets, followed by UV light curing (320–390 nm, Omnicure S1500, Exfo) for 5 minutes at 5 mW/cm^2^. NorHA hydrogel-coated 12 mm cover glasses were then removed from the Sigmacote-treated cover glass, placed in individual wells of a 24 well plate, washed with Phosphate Buffered Saline (PBS) + 1% penicillin/streptomycin, and allowed to swell overnight at 4°C before being used for cell seeding.

#### Mechanical characterization

Cylindrical NorHA hydrogels (5 mm diameter) were fabricated in a syringe barrel and subsequently allowed to swell in PBS in a 24 well plate for 24 hours prior to testing. Unconfined compression measurements were performed at room temperature using a Discovery HR-30 Hybrid Rheometer (TA Instruments) with a constant displacement rate of 20 µm/s. The compressive Young’s modulus was determined from the slope of the stress-strain curve by fitting the linear region between 10% and 20% strain.

### 2.2 Cell culture and hydrogel seeding

#### Endometrial stromal cell maintenance

Primary endometrial stromal cells (PCS-460-010, ATCC; 25-year-old donor) were cultured in fibroblast growth media, containing 10% fetal bovine serum (PCS-201-030 and PCS-201-041, ATCC), and passaged twice prior cryopreservation. All experiments were performed with passage 3 endometrial stromal cells.

#### Hormone preparation

Estrogen (E4389, Millipore Sigma), progesterone (P7556, Millipore Sigma), and cAMP (B7880, Millipore Sigma) were reconstituted in purified water (concentrations of stock solutions were 10 µM, 1mM and 1mM, respectively), sterile filtered, aliquoted, and stored at -80°C. Estrogen and progesterone were diluted 1:1000 and cAMP was diluted 1:20 for cell cultures containing hormones.

#### Endometrial stromal cell seeding

Endometrial stromal cells were thawed and cultured for 48 hours in fibroblast growth media prior seeding at 15,000 cells per cm^2^. Prior to seeding, the hydrogel-coated coverslips were sterilized for 60 min in a germicidal UV box. Sterilization of the hydrogels was followed by a 30 minute media incubation at 37°C (using cell culture media described above). Cells were seeded onto the hydrogels and cultured for 24 hours in fibroblast growth media containing 100 µM GalNAz (N-azidoacetylgalactosamine tetraacylated), as described previously ^[12]^, prior addition of hormones (+Hrm) or without hormones (control, Ctrl). Media was refreshed daily.

### 2.3 Cell and ECM characterizations

#### Nascent ECM labeling

Endometrial stromal cells on hydrogels were washed with PBS, incubated in PBS containing 2 % (w/v) bovine serum albumin and 15 µM AZDye 488 dibenzocyclooctyne (DBCO-488, Vector® Laboratories) for 30 min at 37°C, followed by 5 PBS rinses and fixation in 4% paraformaldehyde for 15 min.

#### Immunofluorescence staining

Cells seeded on hydrogel samples were fixed in Paraformaldehyde (PFA, 4%) for 15 minutes at room temperature, blocked for 1 hour in 2% BSA in PBS (w/v), and incubated in primary antibody (1:200 anti-fibronectin, Sigma F6140 in 2% BSA in PBS (w/v)) overnight at 4°C. Next, the samples were rinsed three times with PBS, followed by incubation in secondary antibody (1:1000 anti-mouse AlexaFluor-568, 1:1000 phalloidin-647, and 1:1000 Hoechst 33342 in PBS) for 1 hour at room temperature. Hydrogels were washed at least 2 times with PBS and stored at 4°C until fluorescent imaging.

#### Enzyme-Linked Immunosorbent Assay (ELISA)

ELISA was performed on endometrial stromal cells cultured atop glass and in GalNAz-free fibroblast growth media containing 2% FBS. Media supernatants from endometrial stromal cell culture were collected after 96 hours without hormones (Ctrl) or with hormones (+Hrm). Prolactin ELISA kit (EHIAPRL, Invitrogen) was used according to vendor protocol.

### 2.4 Imaging and image analysis

Images of endometrial stromal cells stained for nascent ECM, fibronectin, F-Actin and nuclei were acquired on a Leica DMi8 THUNDER widefield microscope with a 25x water immersion objective. All image quantifications were performed using ImageJ^[13]^. Multi-channel image hyperstacks were split into separate channels and analyzed sequentially. First, a CSV header with all measurement labels and several output folders were initialized and created in a user-defined directory for quantification results and mask image storage. Next, each channel was automatically focused using a normal variance-based slice selection helper function *bestSlice (channel window)* – adapted from publicly available code on ImageJ macro wiki ^[14],[15]^ - followed by adaptive background subtraction (*rolling ball radius* determined from noise standard deviation) and contrast enhancement for each slice.

#### Quantification of nuclei

To generate binary masks of the nuclear channel, ImageJ *Huang* method thresholding was applied, followed by *Watershed* segmentation and *AnalyzeParticles/Measure to* quantify nuclei number, area, average size, and percent area coverage. Note that nuclei staining intensities varied with time and analysis thresholds were adjusted accordingly.

#### Quantification of nascent ECM and fibronectin

Analysis was performed across three different slices (*bestSlice* + offsets of −10, 0, +10, constrained within stack bounds). After pre-processing as described above, *Moments* method thresholding was used to generate nascent ECM binary masks. Additional median filtering and morphological cleanup were applied for offset slices.

Binary masks were combined using logical OR operation of *Image Calculator*, followed by quantification of ECM-positive area (in µm^2^) and percent area coverage per image. The same procedure was used for fibronectin quantification.

### 2.5 Statistical analysis

Graphpad Prism 10 was used for all statistical analyses. Each experiment was performed at least three times, and five images were generated for each sample unless specified otherwise. Comparisons between two groups was performed with an unpaired Student’s *t*-test, and comparisons between three groups with Tukey one-way ANOVA. Quantification of nascent ECM and fibronectin area was normalized to the number of nuclei.

## 3. Results and Discussion

### 3.1 Hormones induce decidualization of endometrial stromal cells in vitro

To visualize morphological changes during decidualization *in vitro*, we first cultured endometrial stromal cells without or with progesterone (medroxyprogesterone acetate), estrogen (estradiol), and 3’,5’-cyclic adenosine monophosphate (cAMP) (Figure 2a). Endometrial stromal cells were seeded onto glass coverslips, allowed to attach for 24 hours, and subsequently cultured in control (Ctrl) or hormone/cAMP media (+Hrm) up to an additional 96 hours (Figure 2b). Upon decidualization, decidual stromal cells typically show an increase in cell area with strong F-actin stress fibers, an increase in nuclear area, and are characterized by an increase in prolactin secretion (Figure 2c). Without hormones, cells showed an elongated morphology with little intracellular F-Actin fibers during the 96 hours (Figure 2d and Supplementary Figure 1). Quantification of nuclear area and count of the control group demonstrated no change in nuclear area but an increase in the number of nuclei (Figure 2e), suggesting continued cell proliferation. In contrast, the addition of hormones induced F-Actin stress fiber formation and increased nuclear area (highlighted with white arrows) within 48 hours (Figure 2f), suggesting the onset of decidualization. Similar to the control group, nuclear area showed little change over 48 hours whereas the number of nuclei significantly increased (Figure 2g). There was no additional increase in the number of nuclei in both Ctrl and +Hrm groups when cultured for up to 96 hours (Supplementary Figure 1). In addition, Enzyme-Linked Immunosorbent Assay (ELISA) at 96 hours confirmed the secretion of prolactin (5.4 ± 3.2 pg/mL, n = 3 samples) for +Hrm groups with no detectable secretion in Ctrl groups. These observations confirm previous reports of estrogen and progesterone induced decidualization that is characterized by F-Actin stress fiber formation and an increase in prolactin secretion *in vitro* ^[16]^.

**Figure 2.**
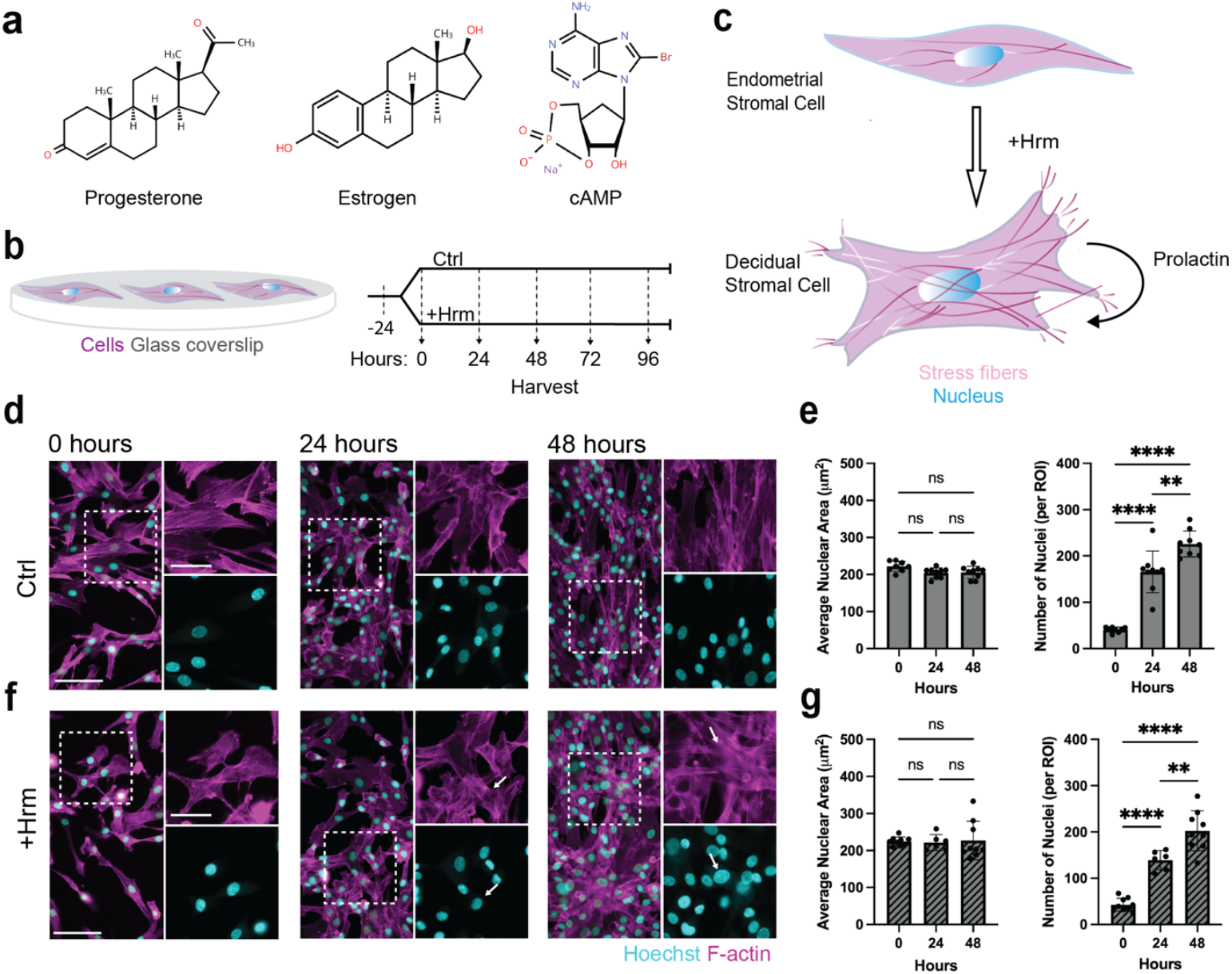
Estrogen and progesterone induce decidualization of endometrial stromal cells. **a** Schematic illustrating the structure of progesterone, estrogen, and 3’,5’-cyclic adenosine monophosphate (cAMP). **b** Timeline of endometrial stromal cell seeding on glass (-24 hours) and culture for 24 hours to reach confluency before an additional culture up to 96 hours without hormones (Ctrl) or with hormones/cAMP (+Hrm). Cells were fixed at 0, 24, 48, 72, and 96 hours for image analysis. **c** Schematic illustrating the change in endometrial stromal cell morphology in response to hormonal treatment leading to the secretion of hormones, such as prolactin. **d** Representative fluorescent images of Phalloidin (F-actin) and Hoechst (nucleus) of endometrial stromal cells cultured for 0, 24, and 48 hours without hormones (Ctrl, scale bars = 100 µm, insets = 55 µm). **e** Quantification of size and number of nuclei of endometrial stromal cells cultured for 0, 24, and 48 hours in control media (n = 3 replicates, 5 images per replicate from one representative experiment, **p≤0.001, **** p≤0.0001, ns = not significant by one-way ANOVA). **f** Representative fluorescent images of Phalloidin (F-actin) and Hoechst (nucleus) of endometrial stromal cells cultured for 0, 24, and 48 hours with hormones (+Hrm, scale bars = 100µm, inset = 55µm). **g** Quantification of size and number of nuclei of endometrial stromal cells cultured for 0, 24, and 48 hours in hormone media (n = 3 replicates, 5 images per replicate from one representative experiment, **p≤0.001, **** p≤0.0001, ns = not significant by one-way ANOVA).

### 3.2 ImageJ enables automated analysis of nascent ECM deposition

After having shown that endometrial stromal cells undergo decidualization, we next aimed to investigate how this process contributes to the spatiotemporal deposition of nascent ECM. We have previously developed sugar-based metabolic labeling to visualize and quantify nascent ECM deposition ^[12]^. Upon addition of an azide-modified sugar (N-azidoacetylgalactosamine-tetraacylated) to standard cell culture media, cells incorporate the modified sugar into glycosylated proteins, such as collagen, and glycoproteins, such as laminin and fibronectin, which are then fluorescently labelled via fluorophore-cyclooctyne (DBCO) mediated click chemistry (Figure 3a). Importantly, nascent ECM labeling is performed prior cell fixation to prevent intracellular labeling. This was followed by staining for nuclei (Hoechst) and the cytoskeleton (F-Actin) for visualization of the cells.

**Figure 3.**
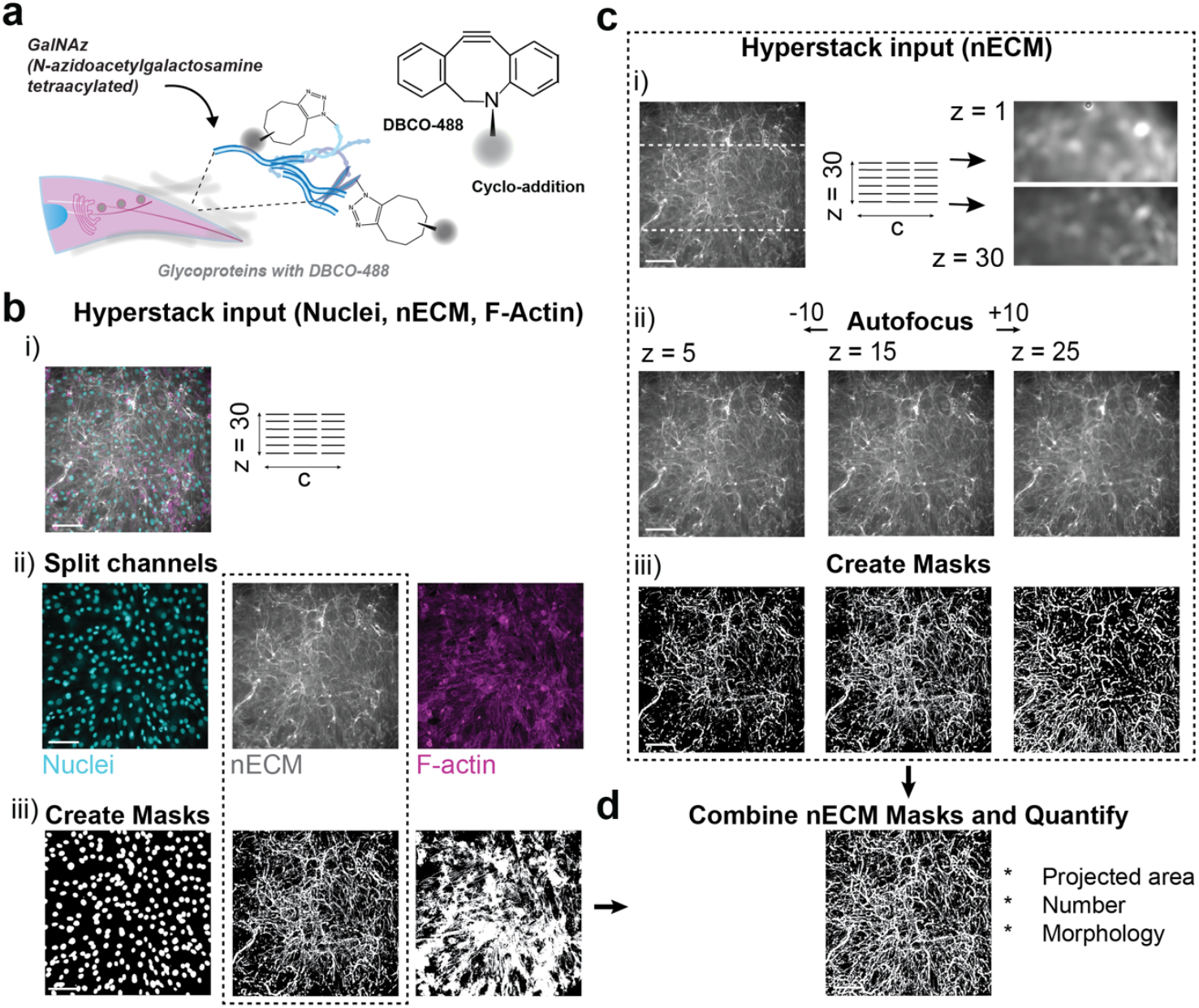
Workflow of automated cell and nascent ECM fluorescent image analyses. **a** Schematic illustrating metabolic labeling of glycosylated nascent ECM via the addition of N-azidoacetylgalactosamine-tetraacylated (Ac4GalNAz) to the media followed by fluorescently tagging glycoproteins using biorthogonal cycloaddition with a DBCO modified fluorophore (DBCO-488). **b** Overview of image analyses workflow with hyperstack images (Nuclei, nascent ECM, and F-Actin, scale bar = 100 µm) that are (i) split into separate channels, followed by (ii) individual threshold and morphological clean-ups to (iii) obtain final images for analyses such as area and number. **c** Example workflow of identifying the focus plane of a fluorescent image with multiple z-planes (scale bar = 100 µm): (i) z-stack images with a step size of 0.63 µm are taken between the upper (z = 1) and lower (z = 30) out-of-focus planes, followed by (ii) identifying the image with the optimal focus plane via autofocus (z = 15) plus one image 10 steps above (z = 25) and one image 10 steps below (z=5). (iii) Each image is then processed into a mask using the threshold function, followed by combining the three images into one compete, pooled mask. **d** Individual masks, both from separate channels and from pooled mask of nascent ECM signal, are quantified to calculate projected area, number of objects and morphology.

To enhance throughput of image analyses, we designed an automated ImageJ-based workflow that enables unbiased and generalizable quantification of multi-channel hyperstack images including fluorescent channels of nuclei, nascent ECM and F-Actin (Figure 3b-i). All images were acquired as hyperstacks that were first split into separate image stacks based on each individual channel (Figure 3b-ii). Next, we applied an automated sequence of autofocus, threshold, and background subtraction of each image prior creating individual masks (Figure 3b-iii). It is important to note that nascent ECM images show high variability in local depth and intensity – a well-known challenge in epi-fluorescent image analysis. For example, within a given image stack (step size = 0.63 µm) the ECM spans many z-stacks (Figure 3c-i). Due to this depth, regions of the nascent ECM are often out of focus when only one plane is used to quantify the ECM. To address this, we first identified the z-stack with optimal focus (e.g., z = 15), followed by collecting a z-plane above (+10 z-slices) and below (-10 z-slices, Figure 3c-ii), and the generation of individual masks for each identified plane (Figure 3c-iii). The combination of these individual masks then enabled the quantification of several features such as projected area, number of objects and morphology (Figure 3d). Comparing the threshold of the nascent ECM from one z-stack to the threshold of the nascent ECM from the three z-stacks described in Figure 3c, the pooled threshold captures much more of the complex ECM architecture than the single threshold alone (Supplementary Figure 2). Taken together, this workflow enables automated and unbiased analysis of complex and heterogeneous fluorescent image stacks.

### 3.3 Hydrogel modulus and hormone exposure regulate nascent ECM deposition

Using metabolic labeling and the newly designed image analysis workflow, we then tested how mechanical properties and decidualization contribute to nascent ECM deposition. To address this question, we cultured endometrial stromal cells for 24 hours prior to adding hormones for an additional 48 hours of culture (Figure 4a). This timeline for hormone exposure was chosen based on the onset of decidualization within 48 hours of hormone exposure when cells are cultured on glass (Figure 2). To test the role of mechanical properties, we first varied the mechanical properties of the underlying substrate using norbornene-modified hyaluronic acid polymers that are fabricated via light-mediated thiol-ene chemistry with different concentrations of the dithiol crosslinker (Figure 4b). Previous work suggests that healthy uterine tissue has an elastic modulus (i.e., Young’s modulus) of approximately 2-5 kPa with higher values in diseased tissue (∼10-15 kPa) ^[8],[9],[17]^. Thus, we fabricated hydrogels with an average elastic modulus of either 5.1 ± 0.187 kPa (1.29 mM dithiol) or 15.8 ± 1.57 kPa (3.24 mM dithiol, Figure 4c). Next, we aimed to confirm that a higher Young’s modulus increases nascent ECM deposition. We cultured endometrial stromal cells in Ac4GalNAz containing media without hormones on softer (5 kPa) and stiffer (15 kPa) gels. Within 48 hours, cells on both 5 and 15 kPa hydrogels formed a continuous monolayer enmeshed in nascent ECM (Figure 4d). While there was little difference in the number of nuclei between 5 kPa and 15 kPa hydrogels, the nascent ECM area per nuclei was significantly higher for cells cultured atop 15 kPa hydrogels (Figure 4e). This is consistent with our previous work showing enhanced nascent ECM deposition by mesenchymal stromal cells when cultured on stiffer hydrogels ^[18]^. Finally, we added hormones to endometrial stromal cells atop 5 kPa hydrogels, which showed a similar nascent ECM structure but nearly a 25% increase in nascent ECM deposition per cell when compared to control groups (Figure 4f). In contrast, addition of hormones to endometrial stromal cells atop 15 kPa hydrogels induced a modest increase of about 11% in nascent ECM deposition per cell (Figure 4g). These data indicate that hormones enhance nascent ECM deposition differently on softer versus stiffer gels, with a more pronounced difference observed on the softer hydrogels. Taken together, these results suggest that both mechanical and biochemical signals regulate ECM deposition during decidualization.

**Figure 4.**
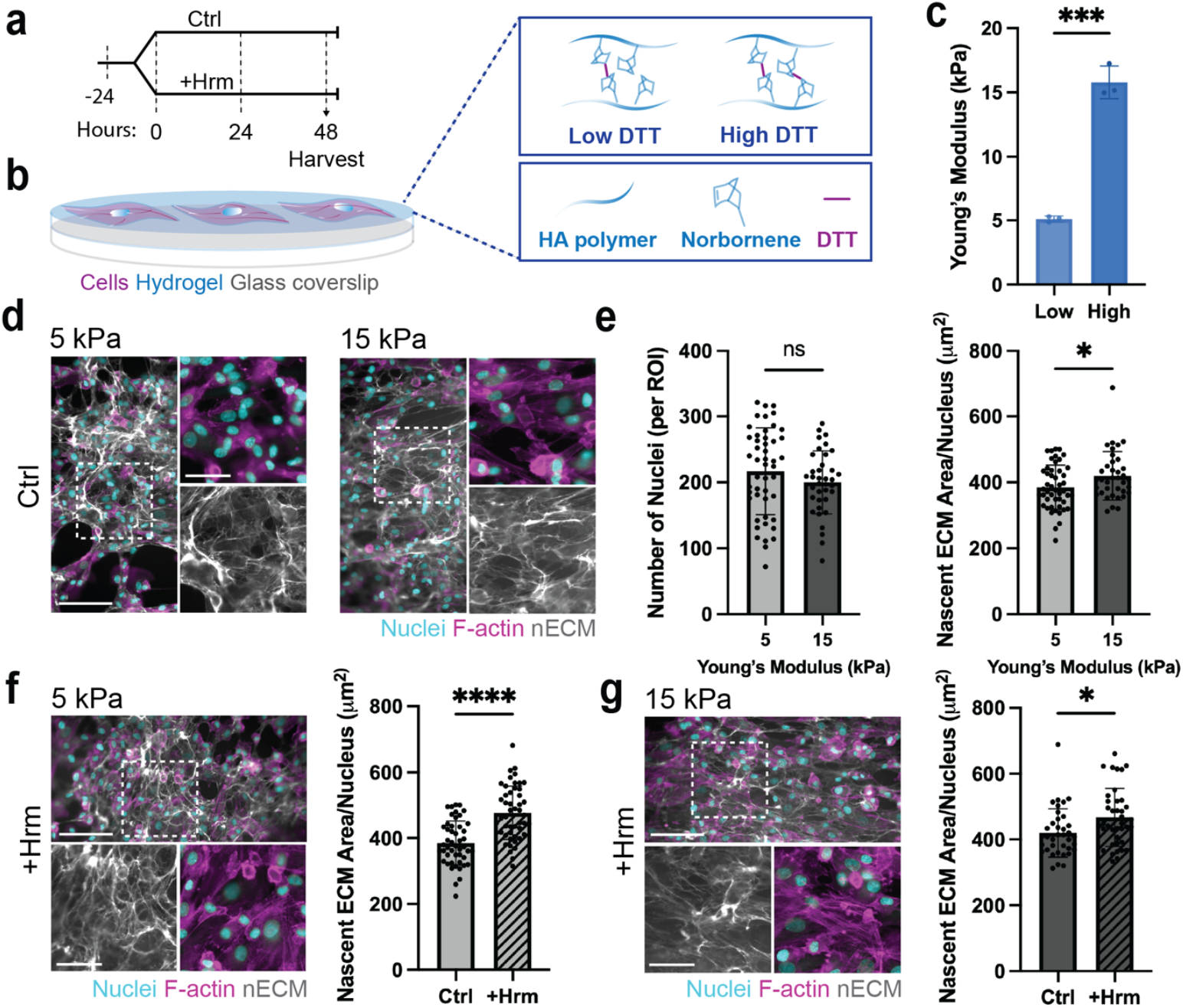
Hormone exposure on softer hydrogels induces a larger increase in nascent ECM deposition. **a** Timeline of endometrial stromal cell seeding: culture for 24 hours to attach to the substrate before additional culture for 48 hours without hormones (Ctrl) or with hormones/cAMP (+Hrm) prior harvesting for image analysis. **b** Schematic illustrating norbornene-modified hyaluronic acid (NorHA) hydrogels with different elastic moduli using 1.29 (low) or 3.24 mM (high) dithiol. **c** Quantification of compressive Young’s modulus of 1.29 mM (low) and 3.24 mM (high) DTT crosslinked NorHA hydrogels (n = 3 hydrogels, ***p=0.0001 by Student’s *t*-test). **d** Representative fluorescent images of endometrial stromal cells cultured for 48 hours atop 5 kPa and 15 kPa hydrogels without hormones (Ctrl, scale bars = 100µm, inset = 55µm). **e** Quantification of number of nuclei and area of nascent ECM deposition, normalized to number of nuclei, of endometrial stromal cells cultured for 48 hours atop 5 kP and 15 kPa hydrogels (5kPa: n = 48 images from 4 independent experiments, 15 kPa: n = 40 images from 3 independent experiments, *p≤0.05, ns = not significant by Students *t*-test). **f** Representative fluorescent images and quantification of number of nuclei and area of nascent ECM, normalized to number of nuclei, deposited by endometrial stromal cells cultured for 48 hours atop 5 kPa hydrogels with hormones (+Hrm, scale bars = 100 µm, insets = 55 µm, n = 48 images from 4 independent experiments, ****p≤0.0001 by Students *t*-test). **g** Representative fluorescent images and quantification of number of nuclei and area of nascent ECM, normalized to number of nuclei, deposited by endometrial stromal cells cultured for 48 hours atop 15 kPa hydrogels with hormones (+Hrm, scale bars = 100 µm, insets = 55 µm, n = 38 images from 3 independent experiments, *p ≤ 0.05 by Students *t*-test).

### 3.4 Hydrogels increase hormone-induced fibronectin deposition

After establishing that both hydrogel modulus and hormonal exposure direct ECM deposition, we next sought to investigate the composition of the newly deposited ECM. To this end, we focused on the glycoprotein fibronectin given its vital role in endometrial tissue health and disease ^[4],[19]^. More specifically, fibronectin has been shown to be highly expressed in the endometrial stroma during the remodeling phase of the menstrual cycle ^[10]^. To explore the relationship between the hormone responsiveness of endometrial stromal cells and fibronectin deposition, we co-stained for the metabolic glycan-labeling and fibronectin via immunofluorescence. On the softer, 5 kPa gels both the control group (no hormones, Ctrl) and hormone treated groups (+Hrm) exhibited strong spatial and structural similarities between fibronectin and nascent ECM staining (Figure 5a). This confirmed that the Ac4GalNAz used for metabolic labeling was incorporated into the glycan chains of the newly deposited ECM. Similar to the nascent ECM readout (Figure 4f) the amount of fibronectin per cell significantly increased with hormone exposure (Figure 5b). The spatial overlap was also observed in the representative images of the control and hormone exposed cells cultured on 15 kPa gels (Figure 5c). Again, much like the nascent ECM per cell, the stiffer gel resulted in a less pronounced upregulation of fibronectin per cell with hormone exposure compared to the softer gels (Figure 5d). Interestingly, although the representative images again showed comparable spatial overlap between the nascent ECM and fibronectin on glass (Figure 5e), when quantified, there was no significant increase in fibronectin with hormone exposure (Figure 5f). Comparing the nascent ECM production and fibronectin across the three substrates, we see reduced ECM and fibronectin per cell in glass compared to the hydrogel substrates, which is particularly obvious in the hormone exposed conditions (Supplementary Figure 3).

**Figure 5.**
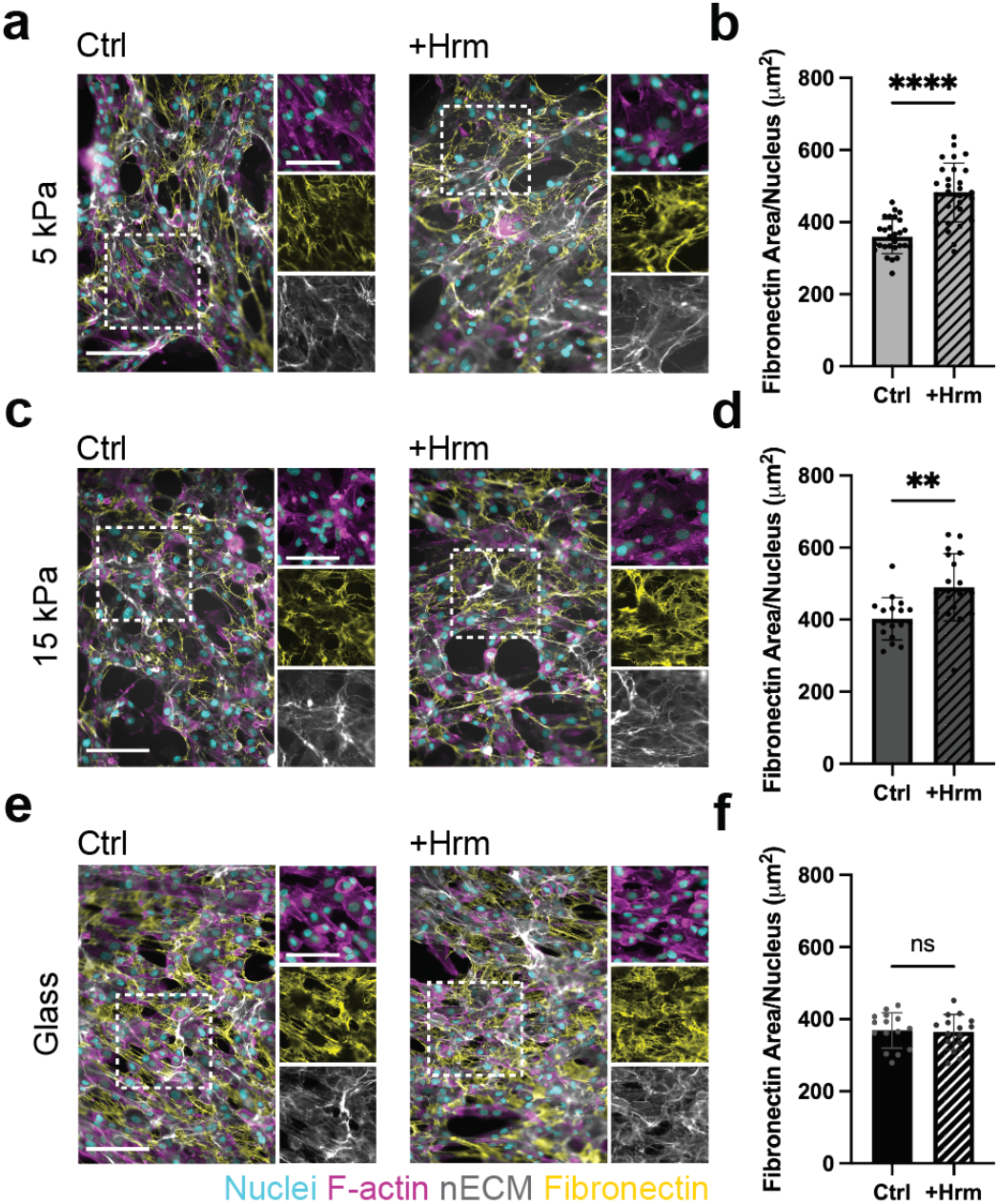
Softer hydrogels promote hormone-induced nascent fibronectin deposition. **a** Representative fluorescent images of cells seeded on 5 kPa gels without (Ctrl) and with hormones (+Hrm), stained with Hoechst (nuclei), Phalloidin (F-actin), DBCO-488 (nascent ECM), and anti-Fibronectin antibody (scale bars = 100 µm, insets = 55 µm). **b** Quantification of area of fibronectin deposited normalized to number of endometrial stromal cells cultured on 5 kPa hydrogels for 48 hours with and without hormones (5 kPa gels: n = 28 images from 3 independent experiments, ****p≤0.0001 by Student’s *t*-test). **c** Representative fluorescent images of cells seeded on 15 kPa gels without and with hormones, stained with Hoechst, Phalloidin, DBCO-488, and anti-Fibronectin antibody (scale bars = 100 µm, insets = 55 µm). **d** Quantification of area of fibronectin deposited normalized to number of endometrial stromal cells cultured on 15 kPa hydrogels for 48 hours with and without hormones (15 kPa gels: n = 19 images from 2 independent experiments, **p ≤ 0.01 by Student’s *t*-test). **e** Representative fluorescent images of cells seeded on glass coverslips without and with hormones, stained with Hoechst, Phalloidin, DBCO-488, and anti-Fibronectin antibody (scale bars = 100 µm, insets = 55 µm). **f** Quantification of area of fibronectin deposited normalized to number of endometrial stromal cells cultured on glass coverslips for 48 hours with and without hormones (glass: n = 15 images from one representative experiment, ns = not significant by Student’s *t*-test).

Our findings suggest that hydrogel culture supports nascent ECM and fibronectin deposition during decidualization and further strengthens the notion that the mechanical properties guide cell function.

## 4. Conclusion

In this study, we engineered hydrogels of varying mechanical properties to investigate the contributions of elastic modulus and hormone exposure to endometrial stromal cell induced nascent ECM deposition. Hydrogels have been widely used for studies of cell mechanosensing and nascent ECM deposition ^[18],[16],[20]^. Yet, a direct relationship between mechanical properties,

nascent ECM, and hormonal stimulation has not been established. By using sugar-based metabolic labeling and estrogen/progesterone addition to endometrial stromal cells, we showed that hormones enhance nascent ECM deposition, which is most effective on soft hydrogels. Our findings further suggest that culturing endometrial stromal cells on glass may suppress their hormone-responsiveness as shown by reduced nascent ECM and fibronectin deposition compared to hydrogels. We anticipate that further modulating the viscous and elastic properties of these hydrogels may enable additional insights into the role of hormones and mechanosensing.

Our work further showed a significant increase in endometrial stromal cell-deposited fibronectin on softer hydrogels. While several other studies have associated fibronectin deposition with fibrotic remodeling or enhanced intracellular contractility, fibronectin is a major component of the healthy endometrial ECM ^[10],[18]^ and thus may play a critical role in endometrial tissue remodeling. It is also important to note that longer time culture may be required to investigate other ECM proteins such as collagens type III and V^[10]^. Additionally, this glycan-based metabolic labeling platform lends itself well to the future study of laminins present in the basement membrane of the glandular and endothelial networks of the endometrium.

Finally, while this study focused on nascent ECM deposition of cells cultured atop hydrogels, 3D culture systems with tunable hydrogel properties or precision-engineered stiffness gradients may be used to further study endometrial stromal cell function and nascent ECM deposition in response to hormone exposure ^[21],[22],[23]^. Taken together, the tunable hydrogel system and metabolic labeling described in this work demonstrate a relationship between tissue modulus and hormone-responsiveness of endometrial stromal cells. These approaches provide a foundation to further investigate the mechanisms of nascent ECM deposition and remodeling in endometrial tissue regeneration and maturation.

## Acknowledgments

This work was partially supported by funding from the NIH (R00-HL151670, R35GM157063 to C.L.), the David and Lucile Packard Foundation (to C.L), and the American Heart Association (to A.R.).

