## Supplement for "Mechanical Cues Regulate Estrogen and Progesterone-Induced Nascent ECM Deposition by Human Endometrial Stromal Cells"

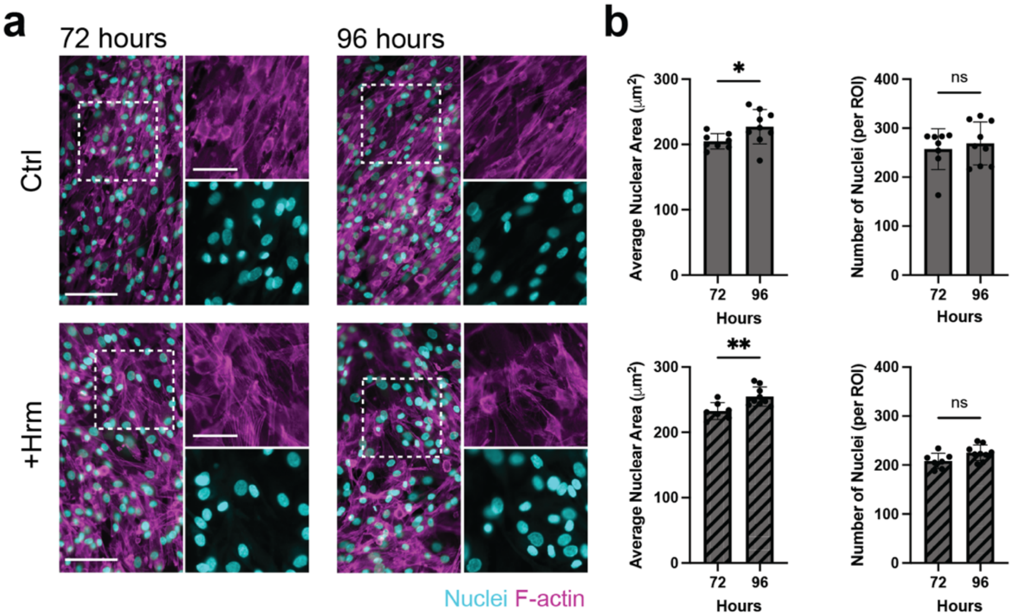


**Supplementary Figure 1. Quantification of nuclear area and number of nuclei on glass**
**a** Representative fluorescent images of Phalloidin (F-actin) and Hoechst (nucleus) of endometrial stromal cells cultured for 72 and 96 h without hormones (Ctrl, scale bars = 100μm in larger image, insets = 55 μm). **b** Quantification of size and number of nuclei of endometrial stromal cells cultured for 0, 24 or 48 h in control media (n = 3 replicates, 5 images per replicate from one representative experiment, * $p \leq 0.05$, ** $p \leq 0.001$, ns = not significant by Student's *t*-test).

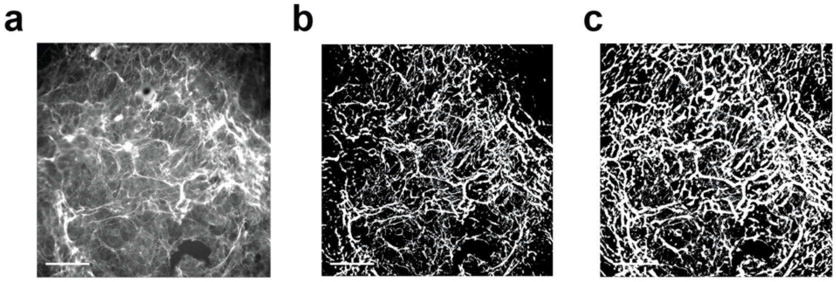


**Supplementary Figure 2. Representative images of nascent ECM, threshold of max projection, and pooled threshold. a** Representative image of nascent ECM max projection. **b** Representative image of threshold of one z-slice exhibiting reduced quantification capacity. **c** Representative image of threshold of three pooled z-slices (as described in Figure 3c) which exhibits complete capture of nuanced details of ECM and improved quantification capacity (scale bars = 100 μm).

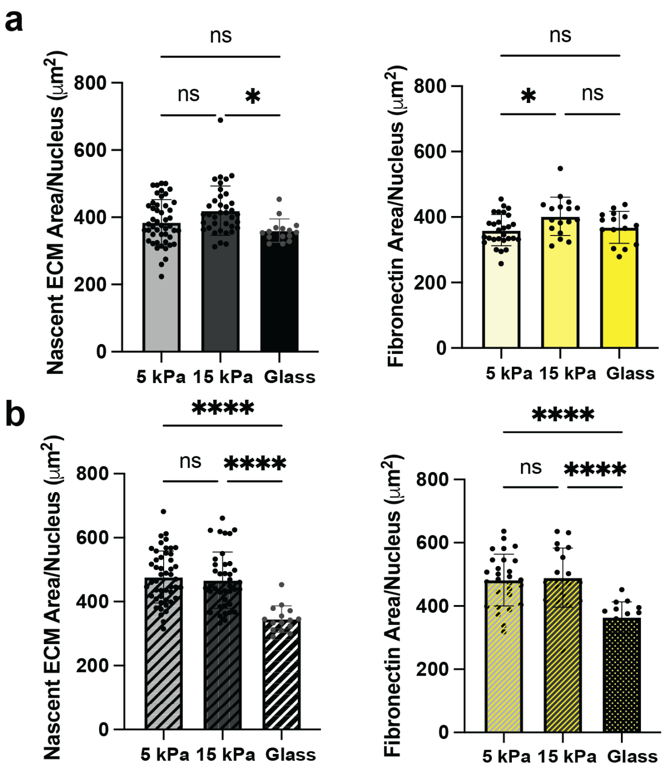


**Supplementary Figure 3. Quantification of nascent ECM and Fibronectin per cell across substrates. a** Quantification of nascent ECM area (grey, $\mu m^2$) and fibronectin area (yellow, $\mu m^2$), normalized to number of nuclei of control (without hormones) group endometrial stromal cells on 5 kPa gel, 15 kPa gel, and glass (5 kPa gels: n = 28 images from 3 independent experiments, 15 kPa gels: n = 18 images from 2 independent experiments, glass: n = 15 images from one representative experiment, *$p \leq 0.01$, ns = not significant by one-way ANOVA). **b** Quantification of nascent ECM area (grey, $\mu m^2$) and fibronectin area (yellow, $\mu m^2$), normalized to number of nuclei of endometrial stromal cells with hormone exposure on 5 kPa gel, 15 kPa gel, and glass (5 kPa gels: n = 28 images from 3 independent experiments, 15 kPa gels: n = 19 images from 2 independent experiments, glass: n = 15 images from one representative experiment, ****$p \leq 0.0001$, ns = not significant, by one-way ANOVA).
